# Is there a sicker sex? Dose relationships modify male-female differences in infection prevalence

**DOI:** 10.1101/2023.05.14.540725

**Authors:** Nathan J. Butterworth, Lindsey Heffernan, Matthew D. Hall

## Abstract

Throughout the animal kingdom there are striking differences in the propensity of one sex or the other to become infected. However, attempts to generalise when we should expect males or females to emerge as the sicker sex have proven challenging. We argue that this is because our current understanding of sex differences in susceptibility is inherently limited, as most inferences have come from field studies (where exposure dose is difficult to quantify), or by measuring infection rates *in vitro* at a limited range of pathogen doses. Without considering how susceptibility changes across a range of pathogen doses (i.e., the dose-susceptibility relationship), we have likely underestimated the scope in which sex differences can arise, reducing our capacity to accurately characterise the ‘sicker’ sex. Here, to expand our scope, we use the *Daphnia magnia* and *Pasteuria ramosa* system to measure infection prevalence across a fifteen thousandfold change in pathogen dose and quantify male and female differences through formal models of environmental transmission. Through this, we reveal that the expression of sex differences in susceptibility is entirely dose-dependent, with males more susceptible at low doses, and females more susceptible at high doses. The scope for male-female differences to emerge is therefore much greater than previously expected – extending to differences in absolute resistance, per-propagule infectivity risks, and the dose-specific behaviour of pathogens. Crucially, none of these components in isolation could define the sicker sex. If we wish to understand the broader patterns underlying whether males or females are the sicker sex, there is a need to apply this expanded scope across the animal kingdom. This will help us understand when and why a sicker sex emerges, and the implications for diseases in nature – where sex ratios and pathogen densities vary drastically.

## INTRODUCTION

Males and females differ in their behaviour, physiology, and genetics – all of which can alter the likelihood and outcomes of infection (Poulin 1996; Zuk and McKean 1996; Sheridan et al. 2000; Zuk 2009; Duneau and Ebert 2012; Gipson and Hall 2016) and the dynamics of disease (Ferrari et al. 2003; Cousineau and Alizon 2014; Úbeda and Jansen 2016; Hall and Mideo 2018; Rogers et al. 2022; Kailing et al 2023). Of all processes, we may expect these sex differences to manifest most consistently in host susceptibility (i.e., the likelihood of infection at exposure), as this is the stage where pathogen encounter and sex differences in immunity first collide (Nunn et al. 2009; Nhamoyebonde and Leslie 2014; Giefing-Kröll et al. 2015). However, the more susceptible sex is in fact highly variable (Sheridan et al. 2000; Kelly et al. 2018) – while in some taxa males seem more susceptible to infection (Poulin 1996; Gray 1998; Adamo et al. 2001; Morton and García-del-Pino 2013; Keiser et al. 2020; Rosso et al. 2020), there are many examples where it is females instead (Seeman and Nahrung 2004; Hillegass et al. 2008; Duneau et al. 2012; Sanchez et al. 2011; Kailing et al. 2023). Thus, while sex differences in susceptibility appear to be widespread, attempts to identify the consistently ‘sicker’ sex have proven challenging (Sheridan et al. 2000; Kelly et al. 2018).

Our understanding of when and why a sicker sex emerges is limited, however, because we have overlooked a major factor underpinning the likelihood of infection – how male and female susceptibility changes with pathogen dose (i.e., the dose-susceptibility relationship; Dwyer et al. 1997; Brunner et al. 2005; Ben-Ami et al. 2010; King et al. 2018; Lunn et al. 2019; Clay et al. 2021; Ng et al. 2022). Susceptibility to infection generally increases proportionally with dose and eventually saturates (following the principal of mass action; Regoes et al. 2003), and as such, differences between sexes that are apparent at low doses can change, or become obscured, at high doses when sex-specific immune responses are overwhelmed. Analysis of this non-linear relationship provides a range of metrics (Figure 1a) which enable us to describe sex-biases in susceptibility beyond simple infection rates. Despite this, most inferences on sex differences have been drawn from studies of wild populations (e.g., Altizer et al. 2004; Rosso et al. 2020; Sweeny et al. 2022) where underlying pathogen dose can, at best, only be approximated (Smith et al. 2005), or from laboratory studies where sex-specific infection rates are often inferred from limited doses (e.g., Gray 1998; Adamo et al. 2001; Duneau et al. 2012; Keiser et al. 2020). In fact, in a recent survey of the pathogen dose-response literature, restricted to one study per host-pathogen combination (Clay et al. 2021), less than five percent of studies included both males and females, and none formally quantified sex differences in the dose-susceptibility relationship. Our current understanding of the ‘sicker’ sex is therefore severely limited by a lack of information on how males and females differ across the full dose-susceptibility relationship (Figure 1).

**Figure 1.**
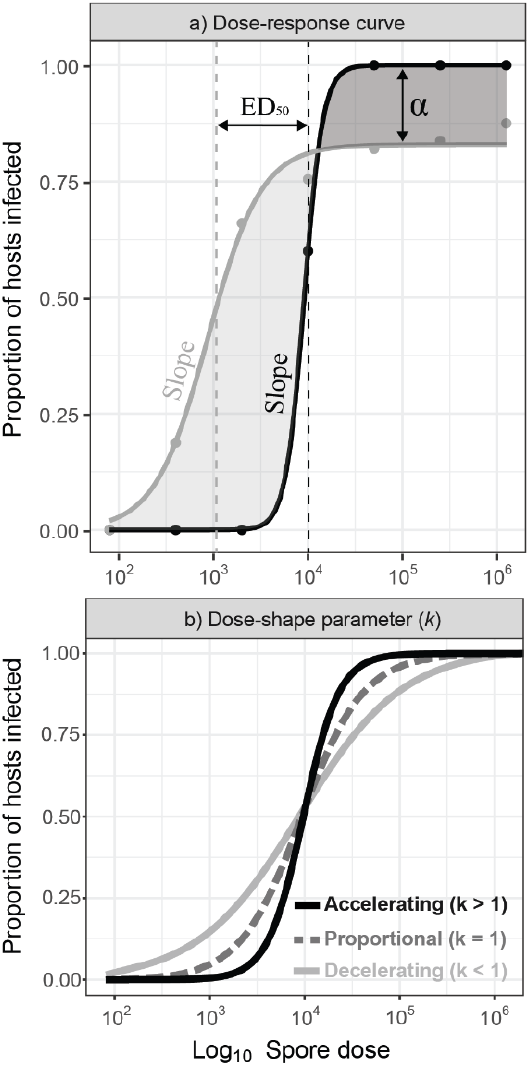
Dose-susceptibility relationships can vary across a range of parameters that can lead to biologically meaningful interpretations **a)** Parameters that can vary between dose-response curves and are quantifiable include: α – the max asymptote (proportion of resistant hosts), ED_50_ the effective dose at which 50% of hosts are infected (roughly equivalent to the mean susceptibility), and the slope of the curve (related to the variance in susceptibility); **b)** A representation of how changes in the dose-shape parameter (*k*, estimated via the parasite synergism/antagonism model of Regoes et al. 2003) alter the dose-response relationship when other parameters are constrained. These include a proportional relationship (*k* = 1; equivalent to mass action), an accelerating relationship (*k* > 1; pathogen synergism), and a decelerating relationship (*k* < 1; pathogen antagonism). All these components may differ between sexes but have yet to be formally analysed.

Another important aspect of susceptibility that has yet to be quantified for males and females is the extent to which the dose-susceptibility relationship diverges from mass action (Figure 1b, Regoes et al. 2003; Clay et al. 2021). As the number of pathogens increases, there can additionally be synergistic (accelerating) or antagonistic (decelerating) dose-response curves (Regoes et al. 2003; Ben-Ami et al. 2008; Clay et al. 2021). Thus, for an individual pathogen particle, it can either become progressively easier to infect hosts as pathogen density increases and immune defences are overwhelmed (synergistic), or alternatively, become progressively harder to infect hosts as intraspecific competition between pathogens begins to interfere (antagonistic, Ebert et al. 2000; Hibbing et al. 2010). This shape parameter (*k*) provides key insight into the density-dependent immune and competitive processes that play out frequently in nature (e.g., Ebert et al. 2000) and is sensitive to various sources of host heterogeneity including physiology and immune activity (Regoes et al. 2003; Ben-Ami et al. 2008). This shape parameter thus provides an entirely overlooked dimension where biologically meaningful sex-differences might arise. By quantifying this alongside the other components of susceptibility (Figure 1a) we have the potential to greatly expand the scope for sex differences to arise, improving our capacity to understand exactly when and why a ‘sicker’ sex emerges.

Here, to formally quantify how sex differences in susceptibility arise across a spectrum of pathogen doses, we use the facultatively parthenogenetic crustacean *Daphnia magna* and its bacterial pathogen *Pasteuria ramosa*. Previous work with this system has pioneered the use of dose-response analyses (Ebert et al. 2000; Regoes et al. 2003; Ben-Ami et al. 2008; Ben-Ami et al. 2010) through which it is well established that the likelihood of infection is tightly linked to pathogen dose, host immunity, physiology, and diet (Ben-Ami et al. 2010). Males have been shown to be more resistant to infection than females at a limited range of doses (Duneau et al. 2012) and the extent of resistance changes with age (Duneau et al. 2012; Gipson and Hall 2018). We therefore expect juveniles of both sexes to share similar patterns of susceptibility, but for sexually mature males to be broadly more resistant to infection than sexually mature females. However, whether this will hold true for a wider range of pathogen doses (i.e., the entire dose-response curve), and when we consider the multivariate view of sex-differences in susceptibility offered by formal models of infection, remains unclear. By disentangling dose, sex, and susceptibility we will provide a formal framework for comparing sex differences in susceptibility within- and between-species.

## METHODS

### Study organisms

*Daphnia magna* Straus is a freshwater crustacean that reproduces via facultative parthenogenesis and can produce genetically identical male and female clones (Ebert 2005). The species is endemic to ponds throughout the Holarctic region (Hebert and Ward 1976; Vanoverbeke and De Meester 1997). *Pasteuria ramosa* Metchnikoff (Green 1974; Ebert et al. 2016) is a natural gram-positive bacterial pathogen of *D. magna*, which invades the host via attachment to the oesophagus and subsequently reproduces within the haemolymph of the infected *Daphnia*, filling the body with transmission spores. In females, infection results in a severe reduction in fecundity and lifespan, and an increase in body size (Clerc et al. 2015; Hall et al. 2019). In contrast, males are naturally smaller, considered more resistant to infection, allow for fewer spores to be produced, and suffer lower virulence (Duneau et al. 2012; Gipson and Hall 2018; Gipson et al. 2022). In both males and females, transmission occurs exclusively horizontally, occurring after the release of spores from a dead host (Hall and Mideo 2018; Aulsebrook et al. 2023).

### Production of experimental animals

This experiment used experimental stocks of *D. magna* of the genotype HU-HO-2, originating from a pond in Kiskunság National Park, Hungary and *P. ramosa* of the genotype C19, originating from an infected female collected from a pond in Gaarzerfeld, Germany (Luijckx et al. 2011). To minimise variation in maternal effects, standardised female *D. magna* were raised individually for three generations in 70 ml jars filled with 50 ml of artificial *Daphnia* medium (ADaM) (Klüttgen et al. 1994; modified by Ebert et al. 1998) and maintained under standard conditions (20°C, 16L:8D).

Genetically identical male and female *D. magna* were produced by exposing the third generation of standardised females to the crustacean juvenile hormone methyl farnesoate (300ug L; product ID: S-0153, Echelon Biosciences, Salt Lake City, UT). Following established protocols, adult females were transferred to 20 mL of hormone after the release of their first clutch (Thompson et al. 2017; Gipson et al. 2022). The second clutch produced after exposure to the hormone, composed of a mixture of male and female offspring, was then collected and immediately returned to normal artificial media. This method of producing both male and female *Daphnia* has no detectable impact on either the lifespan or fecundity of control animals (Thompson et al. 2017). Male and female *Daphnia* offspring were identified by the presence/absence of the sexually dimorphic appendage used for clasping onto females (Ebert 2005).

### Dose-response experimental design

To assess sex differences in susceptibility across a range of doses for both juvenile and mature individuals, males and females to be infected were housed individually in 70 ml jars, filled with 50 ml of artificial media. Animals were fed daily throughout the entire experiment with increasing numbers of algal cells (*Scenedesmus* sp.) – 1 million cells (at 1 day old), 2.5 million cells (at 2-5 days old), 3 million cells (at days 5-6 days old), and 5 million cells (from day 7 onwards). Animals were changed into 50 ml of fresh ADaM every three days throughout the entire experiment. Each treatment consisted of 50 individual replicates for a total of 1,400 experimental animals (2 sexes × 2 ages × 7 doses × 50 replicates).

To infect the animals, *P. ramosa* spores were set up as follows: the highest spore dose of 1,250,000 spores was subsequently diluted seven times by a factor of five to produce the other doses, resulting in seven dose levels of 80, 400, 2000, 10,000, 50,000, 250,000, and 1,250,000 spores. At one-day of age, the ‘juvenile’ cohort of animals were transferred individually into jars containing 20 ml of ADaM and dosed with one of the seven *P. ramosa* spore doses. After 4 days, animals were moved to fresh jars filled with 50 ml of ADaM and then water changed as usual after. At ten-days of age, the ‘mature’ cohort of animals were transferred individually into 20 ml of ADaM and dosed with one of seven *P. ramosa* spore doses. Again, after 4 days, animals were moved to fresh jars filled with 50 ml of ADaM and then water changed as usual after

### Quantifying host susceptibility

Spores of *P. ramosa* infect the haemolymph of *Daphnia* and successful infection induces brownish colouration in both males and females (Ebert 2005; Hall et al. 2019). These phenotypic changes make it easy to identify infection status, which can be determined as early as 14 days after initial exposure (Ben-Ami et al. 2008; Ben-Ami et al. 2010). Experimental animals were thus monitored throughout the experiment until death, or eventually euthanised at 30 days, at which point all individuals had their infection status recorded via their appearance (as per Ben-Ami et al. 2008). In any ambiguous cases (such as animals that died before 14 days post-exposure), animals were crushed, and infection determined via the presence of transmission spores detected by haemocytometer counts under a phase-contrast microscope.

### Statistical analyses

All analyses were conducted in *R* (v. 4.2.1; R Development Core Team 2023) and visualised using the *ggplot2* package (v. 3.4.0; Wickham 2016). A total of 34 animals were removed due to being lost during the experiment. The final dataset therefore contained 1,366 individuals across all seven doses (supplementary material 1). To test how host age and sex contribute to the dose-response relationship, we first modelled the change in the proportion hosts infected with increasing spore dose as a three-parameter log-logistic non-linear model via the package *drc* (v. 3.0-1; Ritz et al. 2016)). This model allows us to partition susceptibility into three components (as per Figure 1a): the ED_50_ (the point at which 50% of hosts are infected), the slope (related to the variance in susceptibility), and the max asymptote (the maximum proportion of infected hosts). For each age, we fit this model twice, once where the parameters were fit to both sexes equally, and then a second time but allowing parameters to vary by sex. We then used an analysis of variance to determine if the sex-specific parameter estimates improved the fit of the constrained model, and thus provide evidence that dose-response relationships varied by sex.

To assess whether the model of environmental transmission differed substantially from mass-action, and whether any changes might depend on the sex of the host, we formally extended our simple approach using the parasite synergism/antagonism model (Equation 1) developed by Regoes et al. (2003), and used previously by Ben-Ami et al. (2008) and Clay et al. (2021). In this model, the proportion of infected hosts at a given dose (*I*_*j*_) is determined by the fraction of resistant hosts (*α*), the per-propagule infectivity of the pathogen (*β*), the pathogen dose (*P*_*j*_), the shape of the dose-response curve (*k*), and the time of exposure (*t*_*exp*_). For *k* = 1, an increase in pathogen dose leads to a proportional increase in *I*_*j*_ (i.e., mass-action). Alternatively, for *k* < 1, an increase in pathogen dose leads to a decelerating increase in *I*_*j*_ (an antagonistic effect of pathogen propagules). For *k* > 1, an increase in pathogen dose leads to an accelerating increase in *I*_*j*_ (a synergistic effect of pathogen propagules) (Regoes et al. 2003).

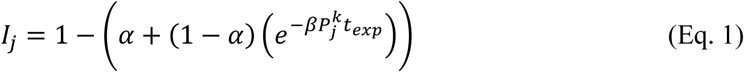

The parasite synergism/antagonism model was implemented in R using the *cmdstanr* (v. 0.5.3; Gabry et al. 2023) implementation of the Stan modelling language (Stan Development Team 2023) and fit to the experimental data. We assumed uniform priors based on a plausible range of values (following King et al. 2018 and Clay et al. 2021). Convergence of the chains was checked visually, and the 90% credible intervals for each parameter were estimated by sampling from the posterior distribution.

Lastly, to determine the contribution of host mortality to observed patterns of susceptibility we also assessed the contribution of increasing spore dose to host mortality (as represented by deaths prior to the 30-day experimental cut off) – using a three-parameter log-logistic model to fit host mortality with increasing spore dose. For each age, we again fit the model twice, once where the parameters were fit to both sexes equally, and then a second time but allowing parameters to vary by sex. We then used an analysis of variance to determine if the sex-specific parameter estimates improved the fit of the constrained model, and thus provide evidence that dose-mortality relationships varied by sex.

## RESULTS

### Sex biases in infection rates are dose dependent and arise in mature animals only

To characterise how sex differences in the relationship between pathogen dose and infection rates might arise, we first estimated each dose-response curve using a three-parameter log-logistic model. For juvenile *Daphnia*, we found no clear statistical support for differences between males and females in their dose-response curves and underlying parameters (*F*_3,8_ = 2.201, *p* = 0.166). Thus, sex differences in susceptibility do not appear to arise in juvenile *Daphnia* at any dose (Figure 2). For mature *Daphnia* however, allowing parameters to vary by sex significantly improved the fit of the dose-response model compared to the model that constrained the curves to be equal for males and females (*F*_3,8_ = 120.190, *p* < 0.001). Sex differences arose in the slope (steeper in females; females = 2.502 ± 0.63, males = 1.536 ± 0.30), max asymptote (higher in females; females = 0.960 ± 0.03, males = 0.842 ± 0.03), and ED_50_ (higher in females; females = 8.135 ± 0.16, male = 6.791 ± 0.17) parameters. As a result, mature males were broadly more susceptible to infection than mature females at low to medium doses, but at high doses a portion of males were unable to be infected by the pathogen (Figure 2).

**Figure 2.**
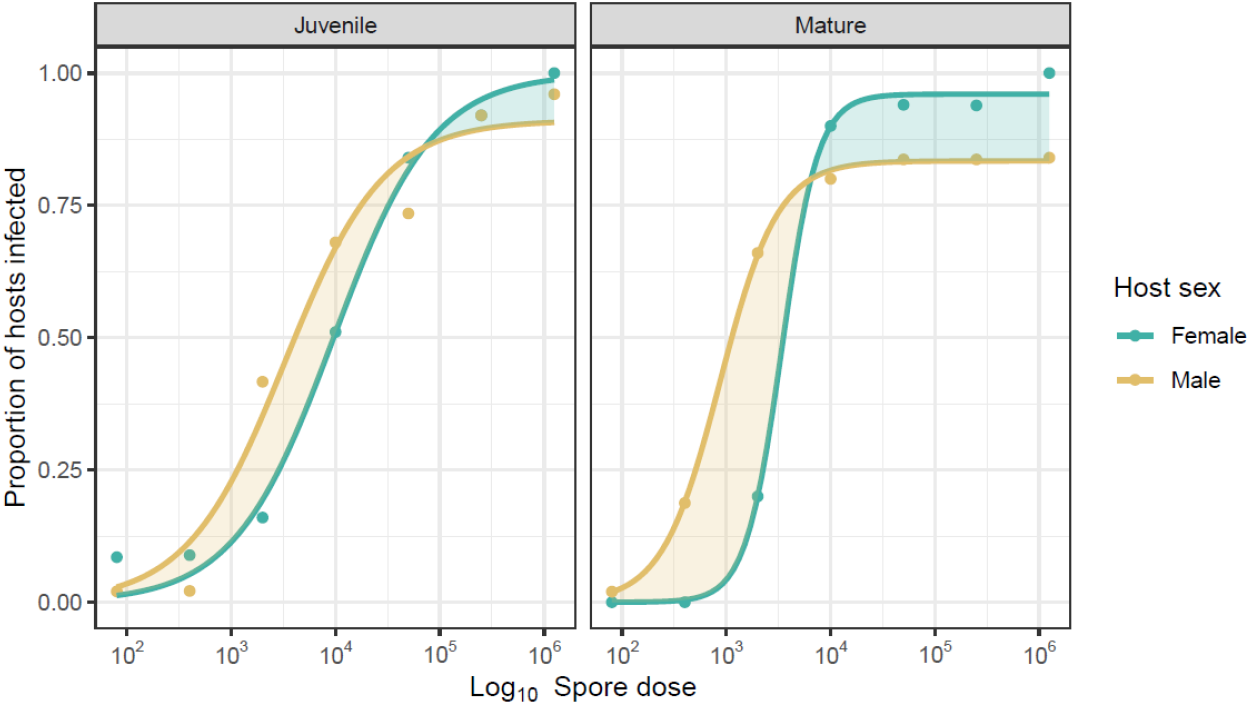
Three-parameter log-logistic models showing the change in proportion of infected *Daphnia magna* (Genotype: HU-HO-2) hosts in relation to the dose of the pathogen *Pasteuria ramosa* (Genotype: C19). Juvenile individuals were exposed to the pathogen when one day old, mature individuals were exposed when ten days old. Yellow shading represents a male bias in infection susceptibility. Green shading represents a female bias in infection susceptibility.

### Ontogenetic changes in environmental transmission occur for females only

To formally quantify these sex differences in terms of the different models of environmental transmission we fit the flexible parasite synergism/antagonism model of infection (Equation 1, Regoes et al. 2003) to our dose-response data. Beginning with the resistant fraction (*α*, Figure 3A), males trended towards a higher *α* value than females, with statistically clear differences emerging (non-overlapping CIs) at maturity. Likewise, for the per-propagule infectivity parameter (*β*, Figure 3B), we again observe that clear differences between the sexes only occur at maturity, with females displaying over a ten-fold lower per-propagule susceptibility than males as an adult. In contrast for males, their per-propagule susceptibility values did not change with ontogeny.

**Figure 3.**
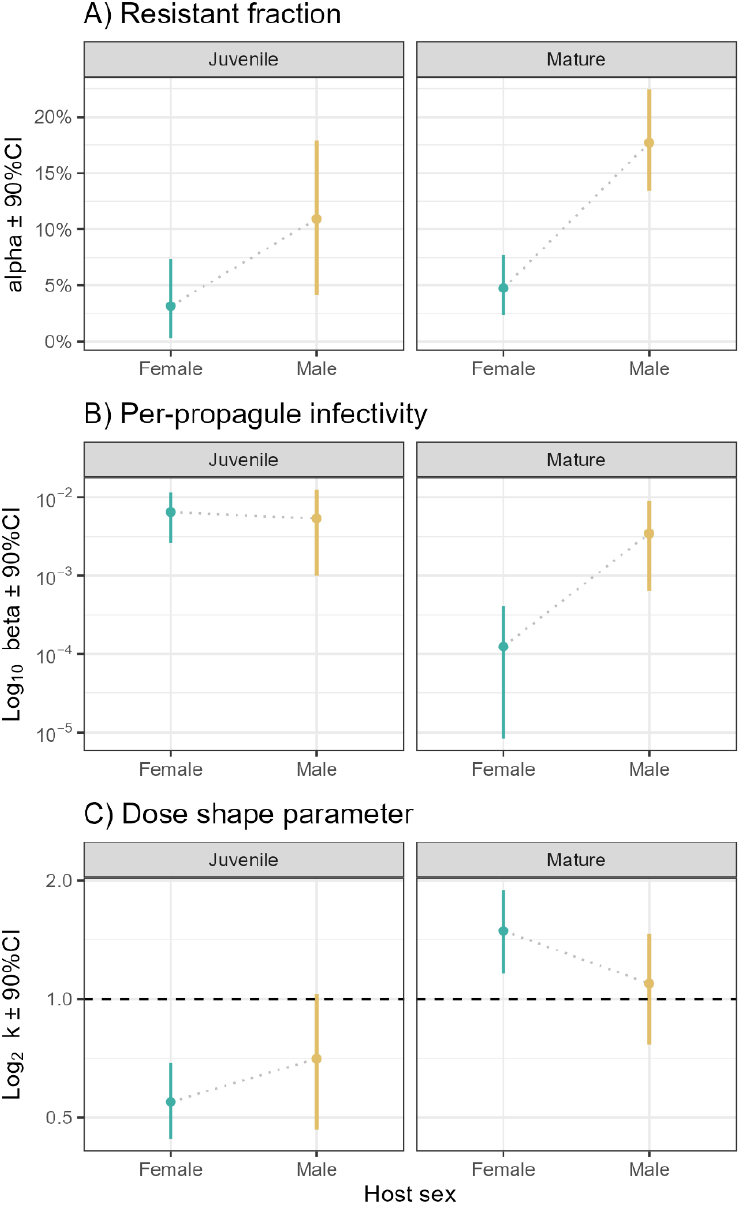
Parameter estimates derived from the parasite synergism/antagonism model of environmental transmission. Shown are the means and 90% credible intervals derived for each age and sex treatment combination. A) *α* corresponds with the fraction of resistant hosts, B) *β* corresponds with per-propagule infectivity of the pathogen, and C) *k* corresponds to the shape of the dose-response relationship.

For the dose-shape parameter (*k*, Figure 3C), we found that for females infection followed an antagonistic relationship as juveniles (*k* ± CIs < 1) and a synergistic relationship as mature adults (*k* ± CIs > 1), indicating a substantial shift in the density-dependent processes shaping pathogen infectivity. For males, although their mean *k* value generally increased across ontogeny, it was largely consistent with a mass-action model at both ages as the CIs indicate no statistically clear difference from a proportional dose-infectivity relationship (*k* = 1). Thus, in line with the previous analysis, the sexes did not differ as juveniles – but diverged at maturity, driven largely by a changing male resistant fraction (*α*), and greater changes in female susceptibility as reflected by both per-propagule infectivity (*β*) and a change in the dose-shape parameter (*k*).

### Mortality is dose dependent and higher in males

A clear finding of our results is that a pathogen is unable to successfully infect and produce mature transmission spores in a fraction of males (the *α* fraction), even at very high exposure doses where infection in females would be completely assured. To determine what differences in male and female life-history may be underlying these patterns, we considered how the mortality of males may be contributing to these results. Shown in Figure 4 is the change in the proportion of exposed hosts that died during the 30-day experiment as partitioned by developmental age and sex. We find that the dose-mortality relationship diverged between the sexes for both juvenile (*F*_3,8_ = 26.302, *p* < 0.001) and mature *Daphnia* (*F*_3,8_ = 67.522, *p* < 0.001). These data contain both infected and uninfected animals, and thus an increased infection rate with dose will naturally lead to dose-dependent increases in mortality. However, beyond this it is clear that this increase in mortality is sex-specific – the relationship between increasing dose and early mortality (< 30 days) is much stronger for males. Males therefore have a much higher likelihood of dying early when infected at high doses, compared to females, hampering the ability of the pathogen to establish a successful infection.

**Figure 4.**
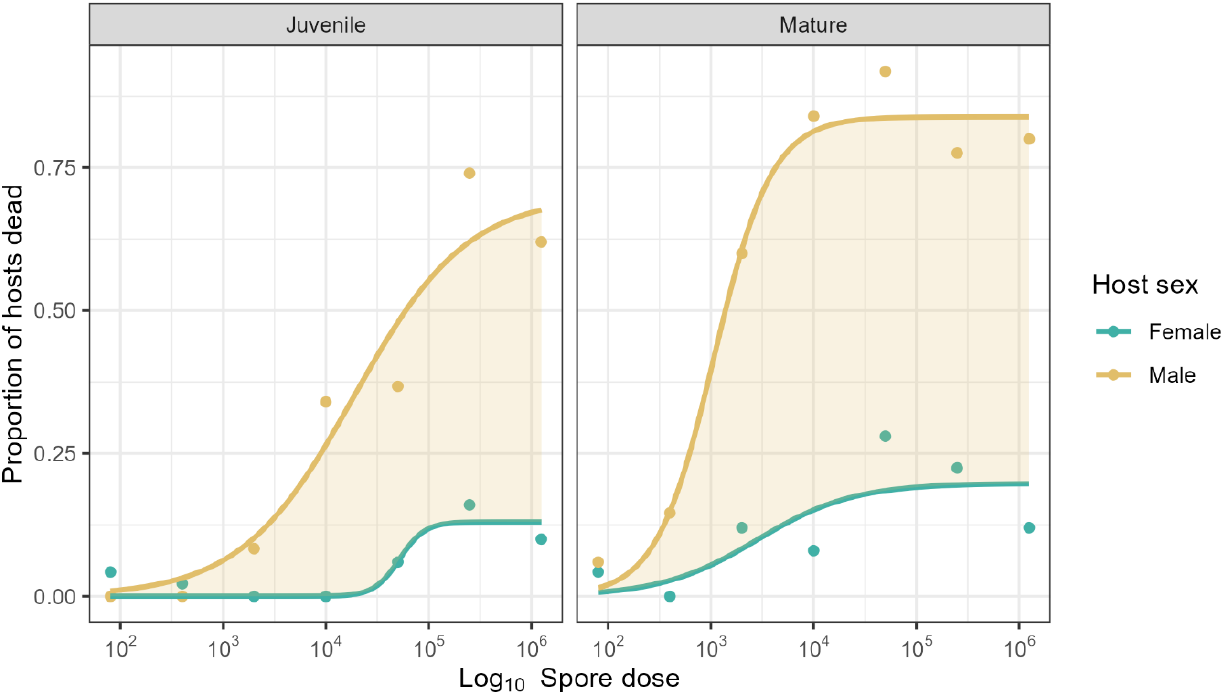
Proportion of male and female hosts exposed to the pathogen that then died during the 30-day experiment, irrespective of their infection status. Juvenile individuals were infected at one day old, mature individuals were infected at ten days old. Curves were modelled with a three-parameter log-logistic function, with the yellow shading highlighting a male bias in mortality rates.

## DISCUSSION

Due to the remarkable variability in the magnitude and direction of sex differences in immune function among species, attempts to generalise the ‘sicker’ sex across the animal kingdom have proven challenging (Sheridan et al. 2000; Kelly et al. 2018). However, studies have yet to assess the sicker sex from the perspective of the dose-susceptibility relationship, so our current understanding is limited by a lack of information on how males and females differ across the entire possible spectrum of susceptibility. Here, we explored how sex differences in susceptibility manifested across a wide range of doses, at two different ages, and formally quantified dose-dependent sex differences with a model of environmental transmission. Our original expectation (based on the prior results of Duneau et al. 2012; but also see Gipson and Hall 2018) was that males of this species would be more resistant to infection as adults. Instead, we found that pathogen dose fundamentally altered the magnitude and direction of sex differences in susceptibility (Figure 2). Based on the metric for which differences in susceptibility are commonly defined, the probability of becoming infected (e.g., Ben-Ami et al. 2010; Clay et al. 2021), males were more susceptible than females at low doses, whereas this pattern was reversed at high doses with females now evidently more susceptible. The expression of the “sicker sex” is thus entirely dose dependent.

Our results reveal how the scope for sex differences to emerge in susceptibility is greater than previously expected and extends beyond simple measure of infection rates. Indeed, every measure of susceptibility that could be derived from a dose-response relationship was found to be sexually dimorphic in adult hosts. For some metrics, for example, males were more susceptible, as they required either a lower pathogen dose for 50% of the cohort to be infected (ED_50_, Figure 2) or their per-spore likelihood of infection was higher (*β*, Figure 3). In contrast, in terms of a resistant fraction (*α*, Figure 2 and 3), females were more susceptible as they always reached 100% infection at high doses, whereas for a portion of exposed males, a successful infection was never established. Thus, no single component of susceptibility could define the ‘sicker’ sex.

The emergence of sex-biases in susceptibility for mature animals alone (Figure 2 and 3) suggests that the emergence of the more susceptible sex is tightly linked to developmental processes (e.g. Adamo et al. 2001; Windsor and Whittington 2010) – as might be predicted based on studies of immune development (Fang et al. 2010; Giefing-Kröll et al. 2015). Underlying this process appears to be a shift in the model of environmental transmission for each sex – as captured by the dose-shape parameter (*k*, Figure 1b) – which reflects how the infectivity of individual propagules changes as pathogen density increases (Regoes et al. 2003). Across both life stages, males maintained a dose-susceptibility (*k*) relationship that could not be distinguished from the mass action model, with the proportion of male hosts becoming infected remaining proportional to dose at all times. Females, however, exhibited a strong shift in *k* from an antagonistic relationship (*k* = 0.54) as juveniles, to a synergistic relationship (*k* = 1.48) as adults (Figure 3C). Thus, a substantial ontogenetic shift in dose-dependent susceptibility occurred in females only, who were harder to infect at high doses when they were juveniles (relatively more ‘resistant’ at high doses), but easier to infect at high doses when they were mature (relatively more ‘susceptible’ at high doses).

This sensitivity of the underlying model of environmental transmission to male and female differences presents an entirely new dimension for understanding the ‘sicker’ sex. Using the same synergistic-antagonistic model of transmission, previous studies have focused on how dose-response relationships are impacted by maternal effects (Ben-Ami et al. 2010), or the species or genotype of host or pathogen (Ben-Ami et al. 2008; Clay et al. 2021). Here, we use the same model to show how throughout development, one sex can be more sensitive to the density-dependent nature of infectivity (females in our case), leading to a three-fold change in the non-linear modifier, *k*. This change, originating intrinsically from the sexes within a single species, is comparable in magnitude to what has been observed among entirely different host-pathogen systems (e.g. Clay et al. 2021) – emphasising the substantial effects of sex differences to the dynamics of infection (see also Úbeda and Jansen 2016; Hall and Mideo 2018; Rosso et al. 2020; Rogers et al. 2022).

Underlying these observed changes in density-dependent infectivity (i.e., the dose-shape parameter, *k*) are likely a range of behavioural and physiological differences that either modify the extent of among-individual heterogeneity in susceptibility (as discussed in Regoes et al. 2003), or shift the extent of intraspecific pathogen competition (Ebert et al. 2000; Regoes et al. 2003) in one sex more than the other. For measures such as ED_50_ and per-spore infectivity (*β*), the lower susceptibility of adult females may reflect a greater investment in immunity than males, which is in line with how immunity and sex differences and roles are viewed from a sexual selection and life-history perspective (Poulin 1996; Zuk and McKean 1996; Adamo et al. 2001; Zuk 2009; Edwards 2012; Metcalf and Graham 2018; Rosso et al. 2020). Physiological differences, however, such as feeding rates (and thus the rate at which males and females encounter the pathogen; see Hall et al. 2007), seem unlikely to be contributing to these components of susceptibility because female *Daphnia* have a higher feeding rates than males in general (Gipson et al. 2022). It is also difficult to explain variation in the resistant fraction (*α*) between sexes by appealing to either immunity or feeding rates – because at the highest doses any immune defences should be overwhelmed (Ebert et al. 2000; Ben-Ami et al. 2010) and encounter rates with the pathogen maximised for both sexes.

Contributing to the dose-dependent shifts in male susceptibility and the overall resistant fraction is instead likely to be differences in the frailty of males and females, both in the presence and absence of infection (Gipson and Hall 2018). Owing to differences in the average lifespan of males and females (Males: 33 days ± 1.9, females: 67 days ± 2.0; Gipson and Hall 2018; Thompson et al. 2017), the pathogen had a much greater chance of killing males compared to females (Figure 4), particularly when infections occur at maturity, as by this point males are already one third through their average lifespan (c.f., only one sixth for females, Gipson and Hall 2018). This increased male frailty manifests as a ‘resistant fraction’, as exposed males died early and before any mature transmission spores could be produced. These spores are needed to successfully generate the secondary infections upon which pathogen fitness depends (Hall and Mideo 2018; Hall et al. 2019). Therefore, while these males may not be functionally resistant, and experience a substantial fitness cost at the individual level, this is still an ecological dead end for the pathogen and a net benefit to the host population – as these pathogens fail to establish in males and are removed from the environment (see analogous arguments regarding patch quality in Nørgaard et al. 2019). Similar processes can be observed in bacterial systems where individual phage-infected bacteria self-sacrifice at an individual cost, but in turn reduce phage spread throughout the population (Lopatina et al. 2020).

Overall, our findings reinforce the notion that males and females fundamentally differ across every measurable aspect of susceptibility, expanding the scope with which sex biases in susceptibility can be considered. Dose-specific responses (e.g. ED_50_), per-propagule infectivity (*β*), resistant fractions (*α*), and even the underlying model of transmission (*k*), are all different ways that a sicker versus more resistant sex can now be defined. Importantly, that the magnitude and direction of sex-differences was specific to each of these components of susceptibility, dose-specific, and that sex differences were not equivalent between life stages (see also Gipson and Hall 2018), helps explain why the more susceptible sex is so hard to characterise – even within a species. If we wish to understand broader patterns underlying whether males or females are the ‘sicker’ sex, there is a need to bring this dose-response framework to studies of sex differences across a much wider variety of taxa. Such data will be crucial for understanding the spread of infectious diseases in nature – where sex ratios, age structure, and pathogen abundance vary enormously.

## Author contributions

Author contributions defined according to the CRediT taxonomy.

Nathan J. Butterworth: Writing – original draft preparation, formal analysis, software, validation, data curation, visualization.

Matthew D. Hall: Conceptualization, methodology, software, formal analysis, writing – reviewing & editing, visualization, supervision, project administration, funding acquisition.

Lindsey Heffernan: Conceptualization, methodology, validation, investigation, project administration.

## Acknowledgements

We thank Kerri Moore, Jared Lush, and other members of the Hall laboratory group for valuable discussion.

## Data availability

Data is available on figshare at 10.26180/22818179

